# Modification of blood-based (IgG) to salivary-based (IgA) ELISA for Inflammatory Bowel Disease Diagnostic: Tumor necrosis factor (TNF) alpha quantification

**DOI:** 10.1101/2022.11.02.514587

**Authors:** Doeun Kim, Anjana Ganesh, Timothy E. Riedel

## Abstract

Approximately 1.6 million people in the United States are struggling with Inflammatory bowel disease. Even though there are a number of diagnostic tools present, including MRI, CT scan, and laboratory tests, the public still lacks access to diagnostic tools due to their expensive costs and needs for the labor of trained phlebotomists. In response, this study focused on modification from blood based enzyme-linked immunoassay (ELISA) to salivary based ELISA in order to expand its accessibility. A 1mL saliva sample was spiked with 1 ± 0.01 μg/mL lyophilized IgG TNFα proteins, and the unspiked saliva was used as a control to test the modified diagnostic. Saliva samples were processed through centrifugation and syringe filtration steps. The change in color between a serum and salivary ELISA kit using either centrifugation or syringe filtration steps was measured by a Color Analysis app that compared red, green and blue values and a microtiter plate reader. The new protocol of salivary-based ELISA lost sensitivity from 31.5pg/mL to 15.6pg/mL of TNFα protein concentration. The best centrifugation method was when a combination of stock saliva and buffer was used before spiking the sample. This means that we can modify the current serum based diagnostic tool to a salivary diagnostic using centrifugation to filter the sample and implement it in developing countries due to its lower cost.

## 1. Introduction

Inflammatory bowel disease (IBD) broadly includes two chronic diseases: Crohn’s disease and ulcerative colitis (*Crohn’s Colitis Foundation*, 2014). Research is still ongoing to examine the causes of IBD, but it is classified as an immunodeficiency disease (Marks et al., 2010) rather than an autoimmune disease as antibodies do not target patients’ own bodies (Amber J. Tresca, 2021). In both diseases, a combination of viruses and antigens trigger the immune system to initiate an inflammatory reaction (*WebMD*, 2019), which leads to malfunction of the gastrointestinal (GI) tract, a set of organs responsible for coordinating the movement and digestion of food (Hornbuckle et al., 2008). The inability of GI organs to absorb nutrients and eliminate waste products cause abdominal pain, diarrhea, anemia, and fatigue (Baumgart et al., 2012; *[National Institute of Diabetes and Digestive and Kidney Diseases](https://www.niddk.nih.gov/health-information/digestive-diseases/crohns-disease/all-content)*). The inflammation damage of ulcerative colitis is limited to the large intestine and the rectum, while Crohn’s disease can damage the entire ileum (small intestine), putting patients at a greater risk of colon and small bowel cancer (Baumgart et al., 2012). Approximately 1.6 million people in the United States are diagnosed with either Crohn’s disease or ulcerative colitis, and the number of new cases of IBD are roughly 70,000 cases per year (Loftus EV et al., 2014). Even though medicines and treatments can help relieve the symptoms, IBD cannot be cured since immunodeficiency states are highly complicated (Amber J. Tresca, 2021). Therefore, early diagnosis and its treatment will be essential to lessen patients’ financial and psychological burdens.

Today, a number of diagnostic tools and therapeutic modalities are available: gastrointestinal endoscopy (Morris et al., 2018), abdominal imaging procedures-X-ray, CT scan, MRI- (*[Mayo Clinic](https://www.mayoclinic.org/diseases-conditions/inflammatory-bowel-disease/diagnosis-treatment/drc-20353320)*), and laboratory tests of biological markers (Nakamura et al., 2003). However, the accessibility of the general public is very low due to expensive costs: cost of imaging procedure range from $500 to $4,000 (*[docpanel](https://www.docpanel.com/blog/post/ct-scan-vs-mri-uses-cost-risks-more)*, 2019) and laboratory tests range from $150 to $600 (Howmuchisit.org, 2018).

Therefore, this study focused on modifying the current blood serum-based enzyme-linked immunosorbent assay (ELISA) to salivary-based ELISA to expand accessibility of exams to the general public. TNFα, which is a protein biomarker that causes inflammation by triggering the production of Interleukin-1 and Interleukin-6, is used as a biomarker for inflammatory bowel disease (Idriss, 2000).

Sandwich ELISA was used to diagnose different levels of biomarkers in the samples. Sandwich ELISA incorporates two antibodies (*[abcam](https://www.abcam.com/protocols/sandwich-elisa-protocol)*) in order to detect quantification levels of the target. An ELISA diagnostic usually receives input from human serum – but collection of these samples requires trained phlebotomists, which can be inaccessible, and expensive to train in developing countries (MacMullan et al, 2020). Current ELISA kits for detecting TNF- Alpha are largely serum based. Other than a serum input, saliva samples can also contain TNF-Alpha and have been detected in periodontal illnesses, so it can be used as an alternative input to the ELISA (Singh, 2014), and are considerably easier to collect, and less expensive, making it easier to use in developing countries. This study investigated quantification of TNFα- a biomarker for inflammatory bowel disease- in two different sample groups: (1) saliva sample spiked with TNF-Alpha and (2) saliva sample with out the protein biomarker. The data was also quantitatively compared to existing blood-serum based ELISA protocol (*[ThermoFisher](https://www.thermofisher.com/elisa/product/TNF-alpha-Human-ELISA-Kit/BMS223-4)*) wi in order to inspect the functionality of the new salivary-based ELISA protocol.

This study tested the following hypothesis: 1) If the sandwich ELISA can detect a difference in salivary samples spiked with TNFα and control, with the level of TNFα ignificantly greater in the spiked sample, and 2) The level output of TNFα from our modified salivary protocol would be similar to output from existing blood-serum based ELISA. Hypothesis 1 is examined by testing a fake saliva- an artificial sample made from active recombinant TNFα proteins- using new salivary-based sandwich ELISA protocol. Hypothesis 2 is examined by comparing the TNFα level with existing Invitrogen™ TNF alpha Human ELISA Kit under the same conditions.

## 2. Materials and Methods

### 2.1. Sample Organism

This study worked with the TNFα antibodies standard, which was purchased from *Thermo Fisher* ([thermofisher](https://www.thermofisher.com/elisa/product/TNF-alpha-Human-ELISA-Kit/KHC3011)). The sample is human blood-serum based, under cytokines and receptors protein family. Genetic base for the sample is homo sapiens- human- with the gene ID of Hu-7124 (Human 7124). The assay will also recognize the recombinant human TNFα, under the same conditions.

### 2.2. Sampling Design

Due to difficulties obtaining real human saliva or blood serum samples of patients with IBD, researchers formulated artificial saliva using lyophilized TNFα proteins.

#### 2.2.1. Saliva Formulation for IBD saliva sample

In order to test the new protocol of salivary-based sandwich ELISA, researchers formulated fake saliva samples through combining lyophilized IgG TNFα proteins to human saliva sample bases (Hettegger et al., 2019). Using these artificial saliva samples, researchers formulated saliva using three different methods: (1) combination of 1 ± 0.01 μL lyophilized IgG TNFα proteins with 1mL saliva base, (2) combination of 150μL culture sample saliva base, 150μL buffer solution with diluent lyophilized IgG TNFα proteins, and (3) combination of 300μL culture sample saliva base with diluent lyophilized IgG TNFα proteins. Each 300μL sample was then used in place of 300 μL standard diluent buffer in the serial dilution procedure for sandwich ELISA.

All saliva samples were stored on ice until handling (about an hour), and centrifuged once before storing. Then samples that are handled within 24 hours were stored at a 4°C fridge, while others were stored at -80°C. In regards to thawing, saliva samples were centrifuged once again for further debris removal (Mandel, 2011). The centrifugation procedure will be further explained below.

#### 2.2.2. Control Sample

Researchers used pure human saliva as negative control of the experiment and it was examined under the same conditions as IBD saliva samples.

#### 2.2.3. Sample Processing

For saliva sample processing, researchers followed the method presented by MacMullan et al.: 1mL of saliva samples were centrifuged under 4°C at 15,000 rpm for 10 minutes duration, and the supernatant was transferred to an empty tube (MacMulan et al., 2020). 300μg of Supernatant was then centrifuged again under 4°C at 15,000 rpm for 8 minutes duration.

### 2.3. Protocol for Data Collection

**Image 1.**
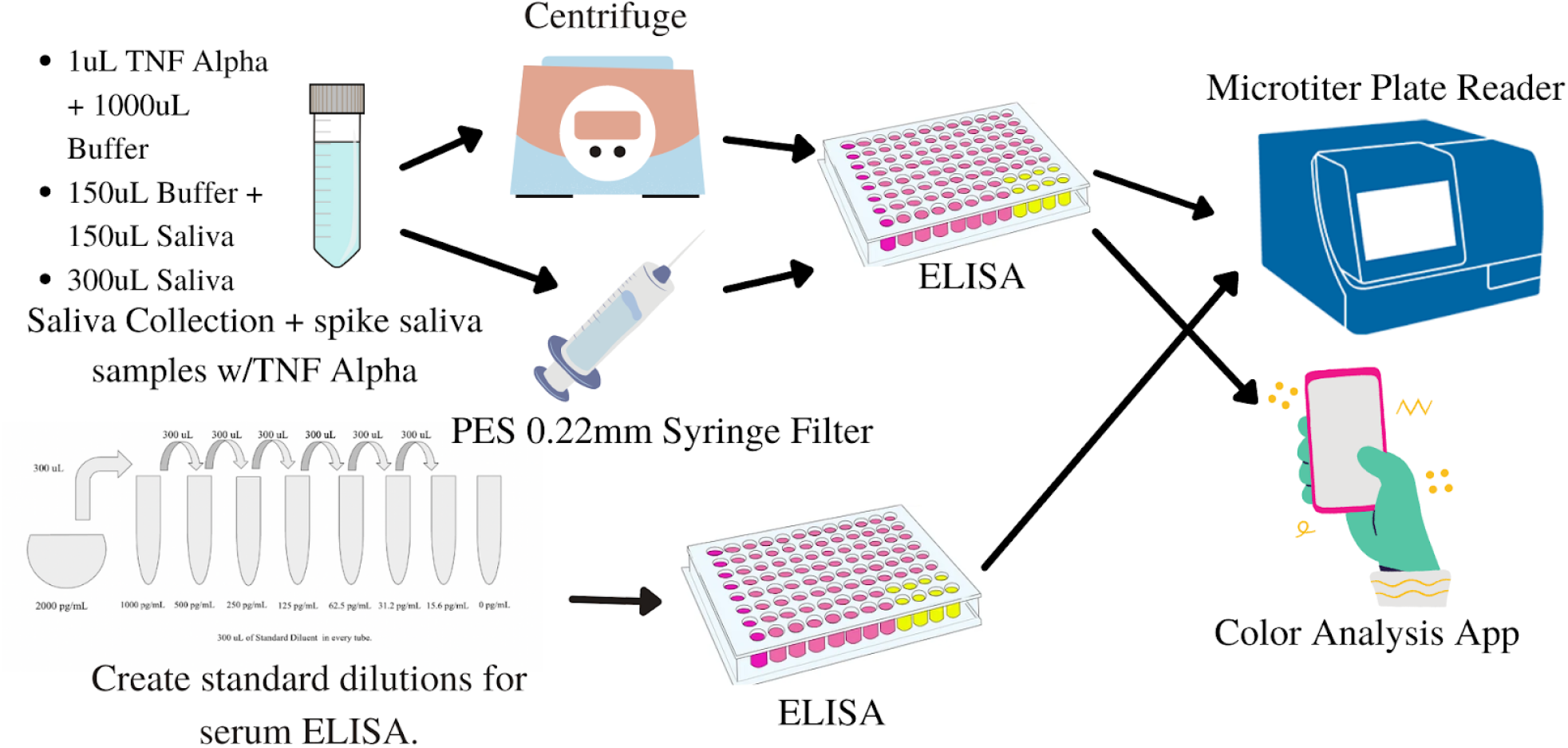
Top row of the flow chart shows the salivary ELISA procedure. Saliva sample was collected and then spiked to 1μL TNF alpha + 1000μL buffer, 150μL Buffer + 150μL Saliva, 300μL Saliva, then either centrifuged or syringe filtered. The bottom row shows the standard dilutions for the ELISA. Both were then read using the microtiter plate reader and color analysis app.

#### 2.3.1. Materials Used for salivary based- ELISA

In order to detect the levels of TNFα in each prepared samples, researchers incorporated the materials list below:

| Materials | Amount |
| --- | --- |
| Hu TNF-a Standard, lyophilized | 0.1% sodium azide |
| Standard Diluent Buffer | 25mL |
| Incubation Buffer | 11mL |
| 96 well plates coated with antibodies | 1 plate |
| Hu TNF-a Biotin Conjugate; contains 8 mM sodium azide | 11mL |
| Wash Buffer Concentrate (25X) | 100mL |
| Chromogen, Tetramethylbenzidine (TMB) | 25mL |
| Stop Solution | 25mL |
| Wash Buffer Concentrate (25X)Streptavidin-HRP, (100X ); contains 1.3 mM thymol | 0.125 mL |
| Streptavidin-HRP Diluent; contains 1.3 mM thymol and 0.05% Proclin™ 300 | 25 m |

The materials listed above are capable of running 6 tests in 96 well plates: 16 wells per test. In addition, supplemental tools and materials including cylinders, plastic tubes, distilled water, and pipettes were used.

#### 2.3.2. Experiment Procedure

##### 1) Sample preparation

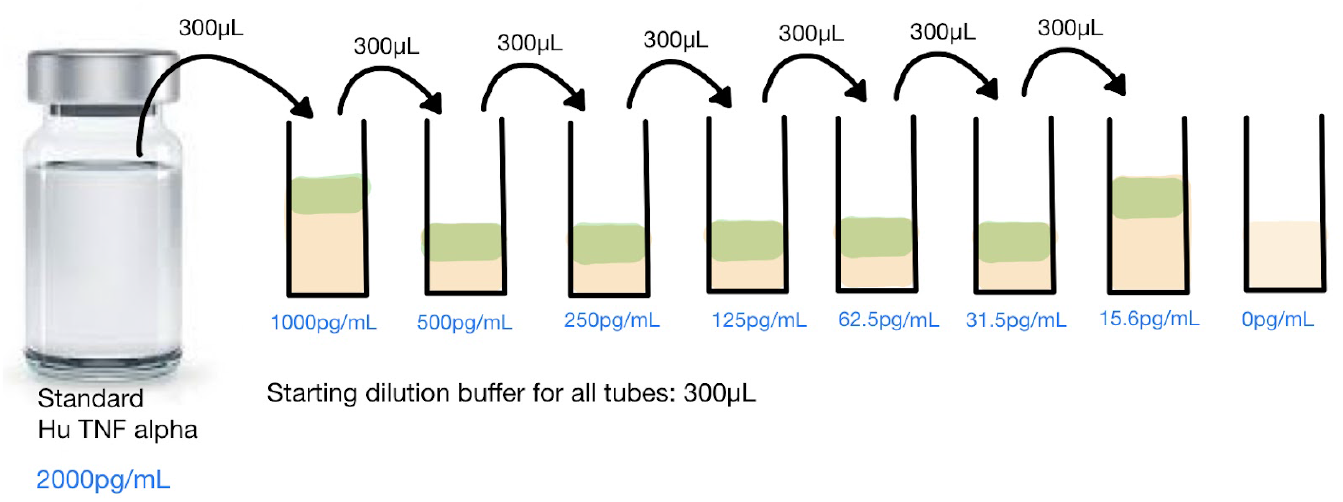

In the preparation process, researchers diluted the reconstituted standard (Hu TNF-a Standard, lyophilized). First, researchers resuspended the standard that has 3.4ng/mL per vial by adding 1.7mL of standard dilution buffer and incubating for 10 minutes. Then after adding 300 μL Standard Diluent buffer was added to each 7 tubes, researchers conducted serial dilution through first adding 300μL of 2000 pg/mL reconstituted standard to the first tube. The serial dilution process is shown in the diagram below:

For the salivary based ELISA, 300 μL of Standard Diluent Buffer was replaced by 300 μL prepared- centrifuged- saliva base (refer back to *2.2.1 and 2.2.3*) to make a complete artificial TNFα saliva sample.

In addition to the diluted standard preparation, 1X Streptavidin HRP solution was also prepared through combining 20μL Streptavidin-HRP solution and 2mL of Streptavidin HRP diluent before 15 minutes of usage.

##### 2) Performing Sandwich ELISA

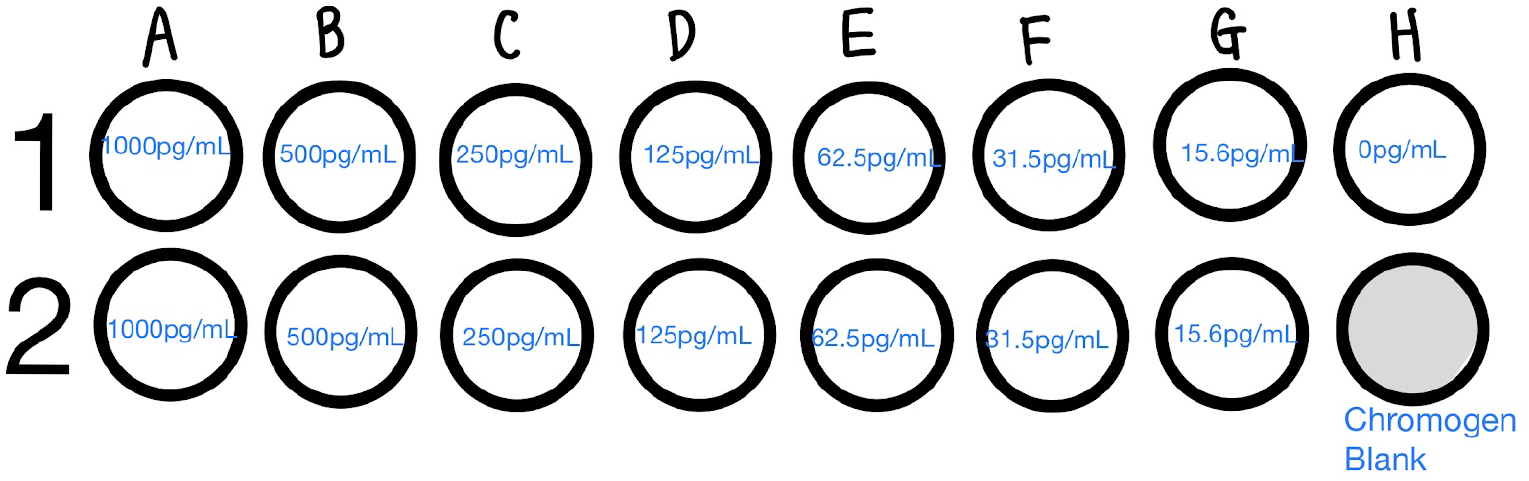

###### 2-1) Antigen binding procedure

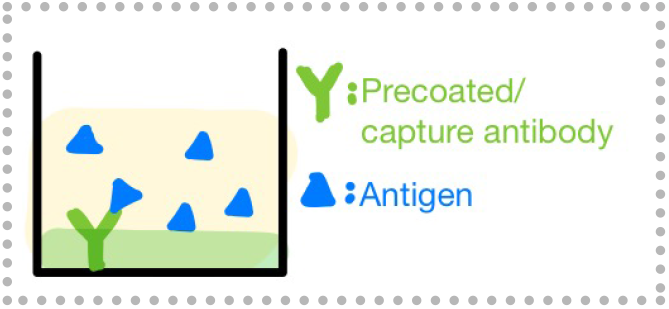

With a prepared solution of diluted standard, researchers added 50 μL of Standard Diluent buffer to each 15 wells and left one well (2H) for the chromogen blank. The 100 μL of prepared sample was also added to each 15 wells except 2H well. This plate was then incubated for 2 hours and washed 4 times with 1X wash buffer.

###### 2-2) Adding Biotin Conjugate

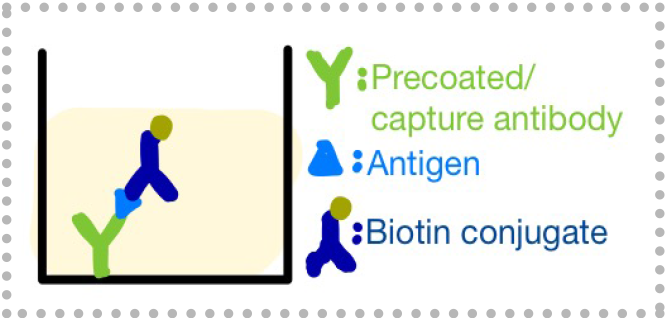

100 μL of TNFα Biotin Conjugate solution was then added to each 15 wells and it was then incubated for 1 hour under room temperature (with the plate sealed with transparent plate cover). After incubation, the plate was washed 4 times with a wash buffer.

###### 2-3) Adding Streptavidin- HRP

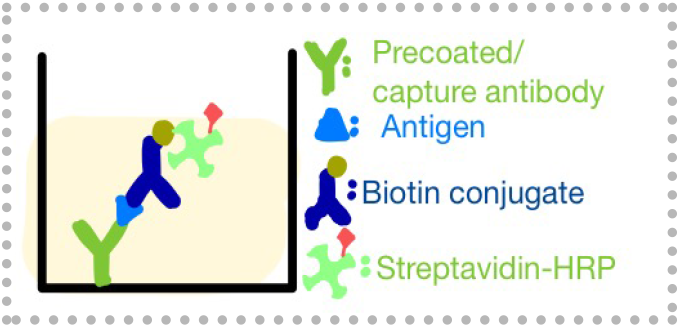

100 μL of 1 X Streptavidin HRP solution was then added to each 15 wells and the plate was incubated for 30 minutes under room temperature. After incubation, the plate was washed 4 times with 1X wash buffer.

###### 2-4) Adding Chromogen

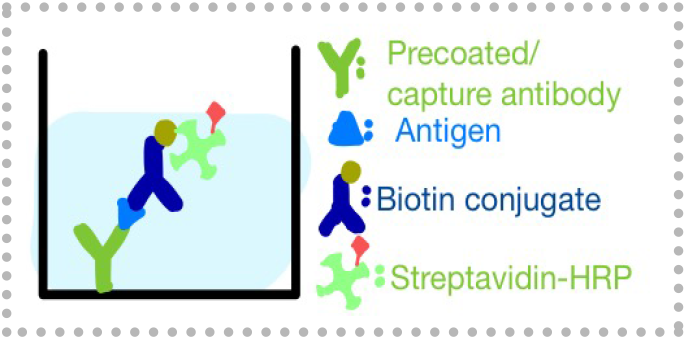

100 μL of room temperature chromogen was then added to every 16 wells and incubated for 30 minutes in the dark. In this process, the solution’s color will change from transparent to blue.

###### 2-5) Adding Stop Solution

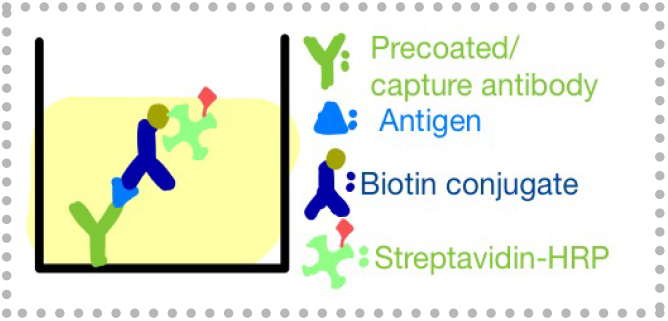

100 μL of stop solution was added to every 16 wells. In this process, the solution’s color will change from blue to yellow according to TNFα concentration.

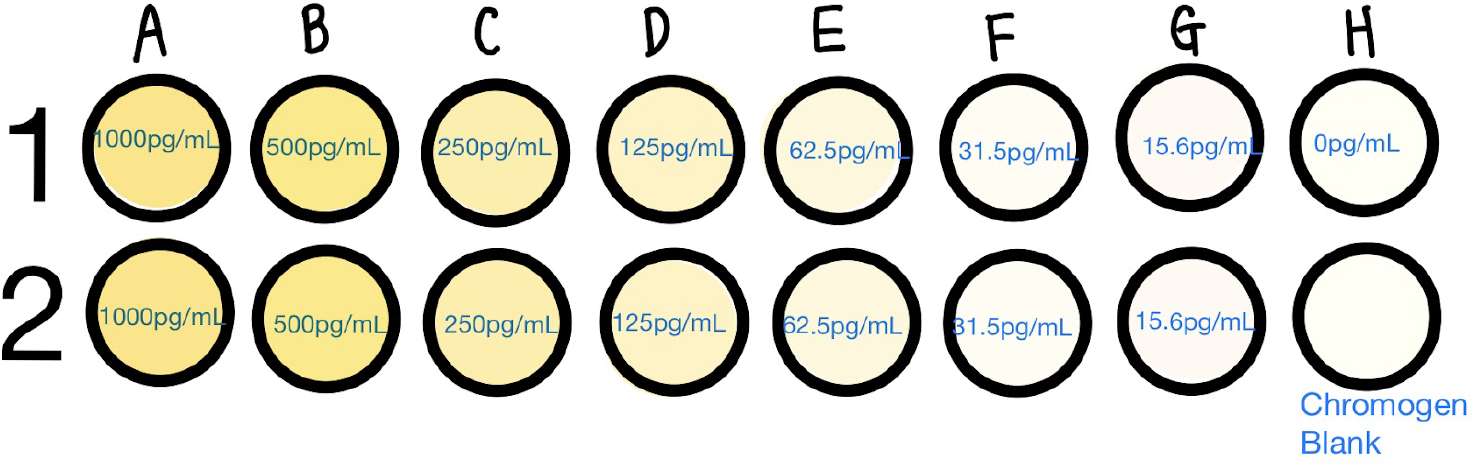

*Note on Washing wells procedure:

In steps (2-1), (2-2), and (2-3), researchers wash the well plate 4 times using 1X Wash Buffer solution. In these procedures, researchers used pipette to place solution to each plate instead of using a squirt bottle in order to prevent buffer forming.

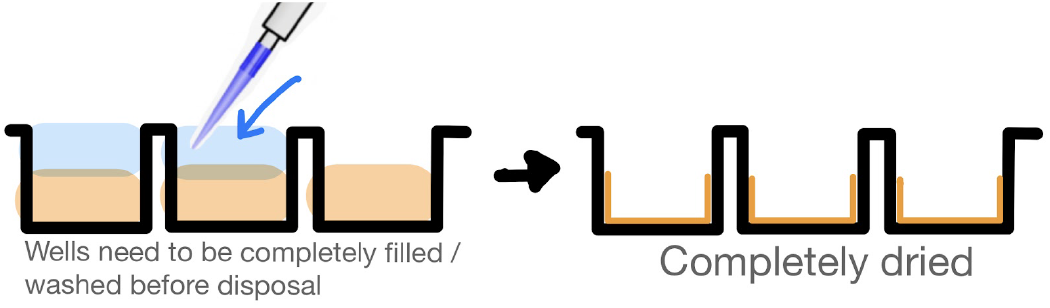

### 2.4. Data Analysis

#### 2.4.1. Qualitative analysis

ELISA plate result is readily detectable with human eyes qualitatively through looking at the color gradient. The more dense and opaque the color is, the more TNFα biomarkers that are detected.

#### 2.4.2. Qualitative analysis using Color quantifier App

For more advanced and easier qualitative analysis of well plates, researchers created an app that makes at-home diagnostics possible: Color Quantifier App. In the following app, users can upload the photo of the 96 well plate, and the computer will analyze the color gradients for users. (Link: https://doeun-kim27.github.io/color-quantifier/).

#### 2.4.3. Statistical, Computational analysis

While the ELISA plate result is easily readable without use of a machine, this study used Spectramax Pro software to further analyze the 96 well plate to quantify the level of TNFα that is detected. For the data analysis process, synergy HT hardware and gen5 software was used to read the well plate. In this procedure, the plate was read under the absorbance at 450nm and the data was analyzed by generating a standard curve of each wells’ optical activity. The optical density datas were then collected and compared for each 8 different sample groups. The result should look similar to the standard curve data below (*[Thermofisher](https://assets.thermofisher.com/TFS-Assets/LSG/manuals/MAN0003931_KHC3011_HumanTNFalpha_ELISA_PI.pdf)*).

| Human TNF-a (pg/mL) | Optical Density (450 nm) |
| --- | --- |
| 1000 | 3.38 |
| 500 | 1.97 |
| 250 | 1.18 |
| 125 | 0.75 |
| 62.5 | 0.38 |
| 31.2 | 0.22 |
| 15.6 | 0.15 |
| 0 | 0.08 |

However, this procedure is not required for the final at-home ELISA kit. This analysis procedure was used only to more accurately examine the newly made ELISA protocol and compare it with existing Invitrogen™ TNF alpha Human ELISA Kit.

## 3. Results

### 3.1. Qualitative Laboratory Results and Data Analysis

#### 3.1.1. Sensitivity of blood based ELISA

Prior to conducting salivary-based ELISA, researchers ran the blood-based ELISA, precisely following the existing protocol of ELISA kit Thermofisher (*[Thermofisher](https://assets.thermofisher.com/TFS-Assets/LSG/manuals/MAN0003931_KHC3011_HumanTNFalpha_ELISA_PI.pdf)*). Each column contains 8 different dilution samples: concentrations are 1000pg/mL, 500pg/mL, 250pg/mL, 125pg/mL, 62.5pg/mL, 31.2pg/mL, 15.6 pg/mL, and 0pg/mL, accordingly from top to bottom. TNFα proteins were detected in every concentration from 1000pg/mL to 16.5pg/mL, changing the solution color to yellow. Higher concentration of TNFα protein showed greater yellow color gradient. For the control- a well with 0pg/mL TNFα- and chromogen blank, both wells showed clear color. For the duplicate samples of the blood based ELISA, no significant difference was observed. The result Image is shown in Image 1.

#### 3.1.2. Sensitivity of salivary based ELISA

Under the same conditions with blood based ELISA, researchers ran the salivary based ELISA in three different methods, differing by artificial saliva formation procedures: (1) combination of 1 ± 0.01 μL lyophilized IgG TNFα proteins with 1mL saliva base, (2) combination of 150μL culture sample saliva base, 150μL buffer solution with diluent lyophilized IgG TNFα proteins, and (3) combination of 300μL culture sample saliva base with diluent lyophilized IgG TNFα proteins. For Column 2, top 4 wells, which are artificial samples containing TNFα proteins showed visible color change, while the bottom 4 wells- control saliva samples showed no color changes. For Column 3 and 4, they followed same concentration gradient dilution from standard ELISA: 1000pg/mL, 500pg/mL, 250pg/mL, 125pg/mL, 62.5pg/mL, 31.2pg/mL, 15.6 pg/mL, and 0pg/mL, respectively from top to bottom. The color change of the solution was visible up to 31.2pg/mL. Salivary-based ELISA lost 50% of sensitivity from 31.2pg/mL to 15.6pg/mL, but the disease proteins are still detectable up to 31.pg/mL concentration. The result Image is shown in Image 2.

**Img. 1-.**
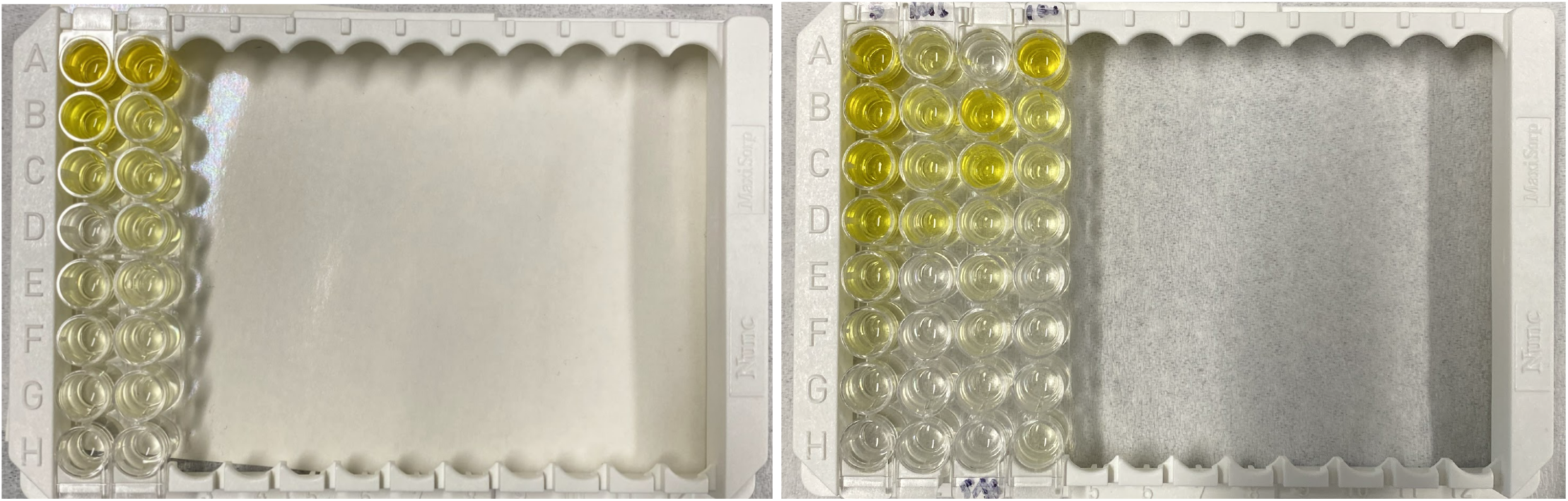
Photo of blood-based ELISA result: The experiment was conducted as a duplicate: samples in the first column and second column are identical except row H. Well H1 is an assay control and well H2 is a chromogen blank. The color gradient difference is clearly visible: As the TNFα concentration increases from bottom to top, the color gradient also increases. This photo was taken right after adding the stop solution-step 5 of sandwich ELISA.

**Img. 2-.** Photo of salivary-based ELISA result: The first column of the plate is ELISA standard: standard blood serum based ELISA result. Then column 2,3, and 4 is salivary based ELISA with three different methods: (Column 2) combination of 1 ± 0.01 μL lyophilized IgG TNFα proteins with 1mL saliva base, (Column 3) combination of 150μL culture sample saliva base, 150μL buffer solution with diluent lyophilized IgG TNFα proteins, and (Column 4) combination of 300μL culture sample saliva base with diluent lyophilized IgG TNFα proteins. This photo was taken right after adding the stop solution-step 5 of sandwich ELISA.

*\*Note on Experimental Errors:*

During the lab procedures, researchers left the A3 well out for the salivary ELISA plate, which is the reason why A3 well is not showing the color change although it is supposed to show greater color gradient than B3. Therefore, data from A3 was eliminated for the future analysis procedure.

#### 3.1.3. Qualitative Analysis: Sensitivity comparison between blood based ELISA and salivary based ELISA

The sensitivity of the new protocol decreased 50%. Color gradient was detectable up to 31.2pg/mL, but undetectable in 16.5pg/mL well.

#### 3.1.4. Qualitative Analysis: Sensitivity comparison between different methods of artificial saliva making procedure

Second method- combination of 50μL culture sample saliva base with diluent lyophilized IgG TNFα proteins- worked much more efficient in comparison to the third method- combination of 300μL culture sample saliva base with diluent lyophilized IgG TNFα proteins. The change in color was detectable as low as 15.6pg/mL for the Column 3 (Method2), while change in color was detectable as low as 62.5pg/mL for the Column 4 (Method3).

### 3.2. Quantitative Analysis Results

**Img. 3-.**
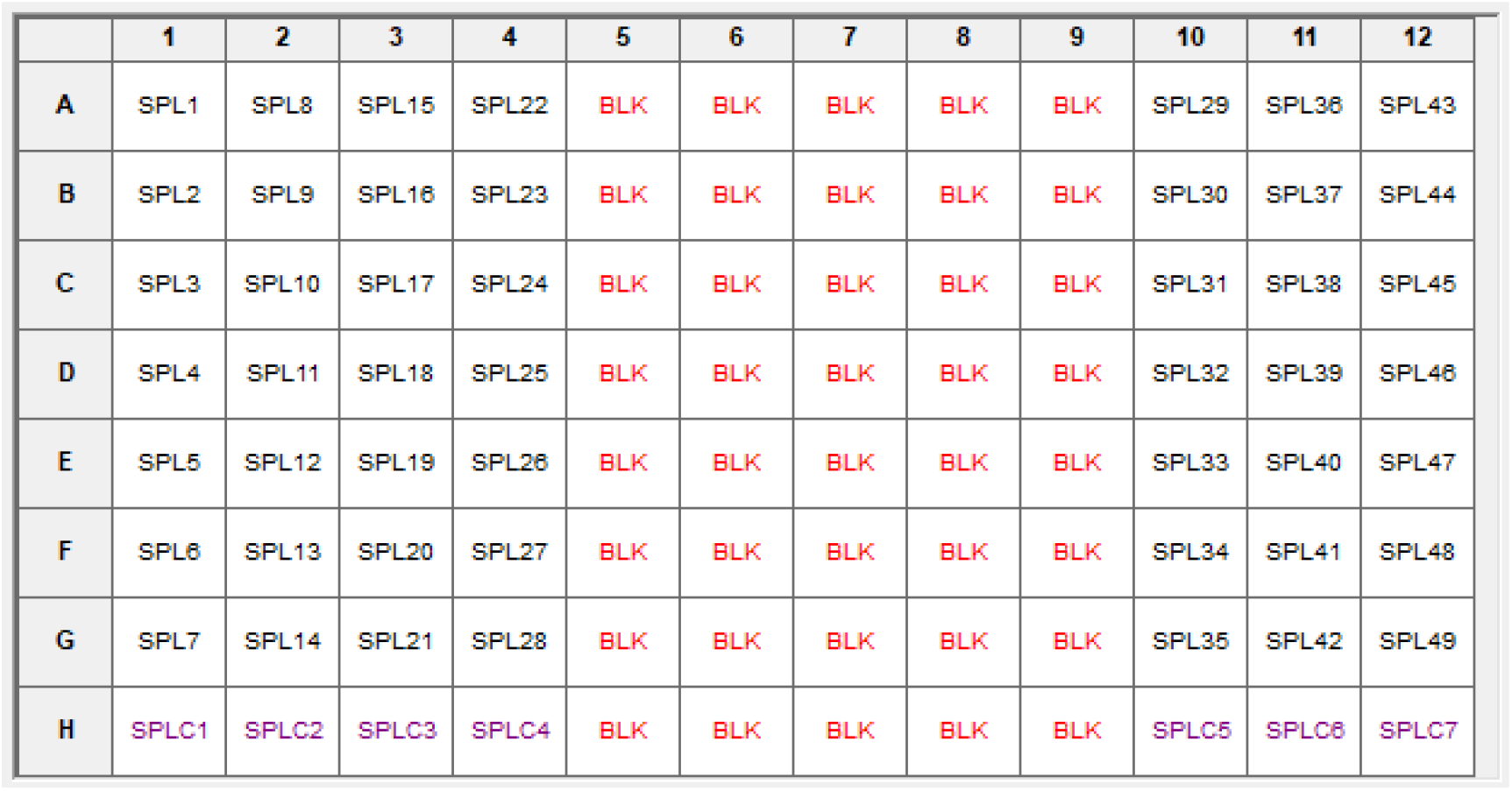
Image of how quantitative analysis on gen5 software was set-up. Chromogen blank/ empty samples are color coded as red. Assay controls/ sample controls are color coded as purple.

#### 3.2.1. Quantitative Analysis sample

Both of the test plates were placed into the -80°C freezer after each ELISA run. A plate of the first blood serum based ELISA was analyzed after 3 weeks, and a plate with saliva based ELISA with serum ELISA standard was analyzed after 10 days. Due to frequent freezing-thawing procedure of the plate1- blood serum based ELISA-, most of the proteins were degenerated and therefore was unable to proceed the analysis procedure under spectramax. For the spectramax analysis, only a plate with saliva based ELISA with serum ELISA standard was analyzed.

#### 3.2.2. Determination of wavelength of analysis

Before proceeding with the TNFα quantification analysis, researchers read the plate under different wavelengths in order to determine the best wavelength of analysis. According to the data- refer to Table 3 and Graph 1- light absorption level ranged from 0.061Au to 0.072Au under 300 nm, 0.03Au to 0.548Au under 450nm, and -0.004Au to 0.033Au under 600 nm. A slope of light absorbance level vs. different concentration levels of TNFα proteins under 450 nm showed greatest magnitude of the slope. Meanwhile, the slope under 300 nm and 600 nm was very close to 0. Therefore, researchers concluded 450nm is the most ideal wavelength for ELISA plate analysis as it gives the most variety of light absorbance levels.

**Graph. 1-.**
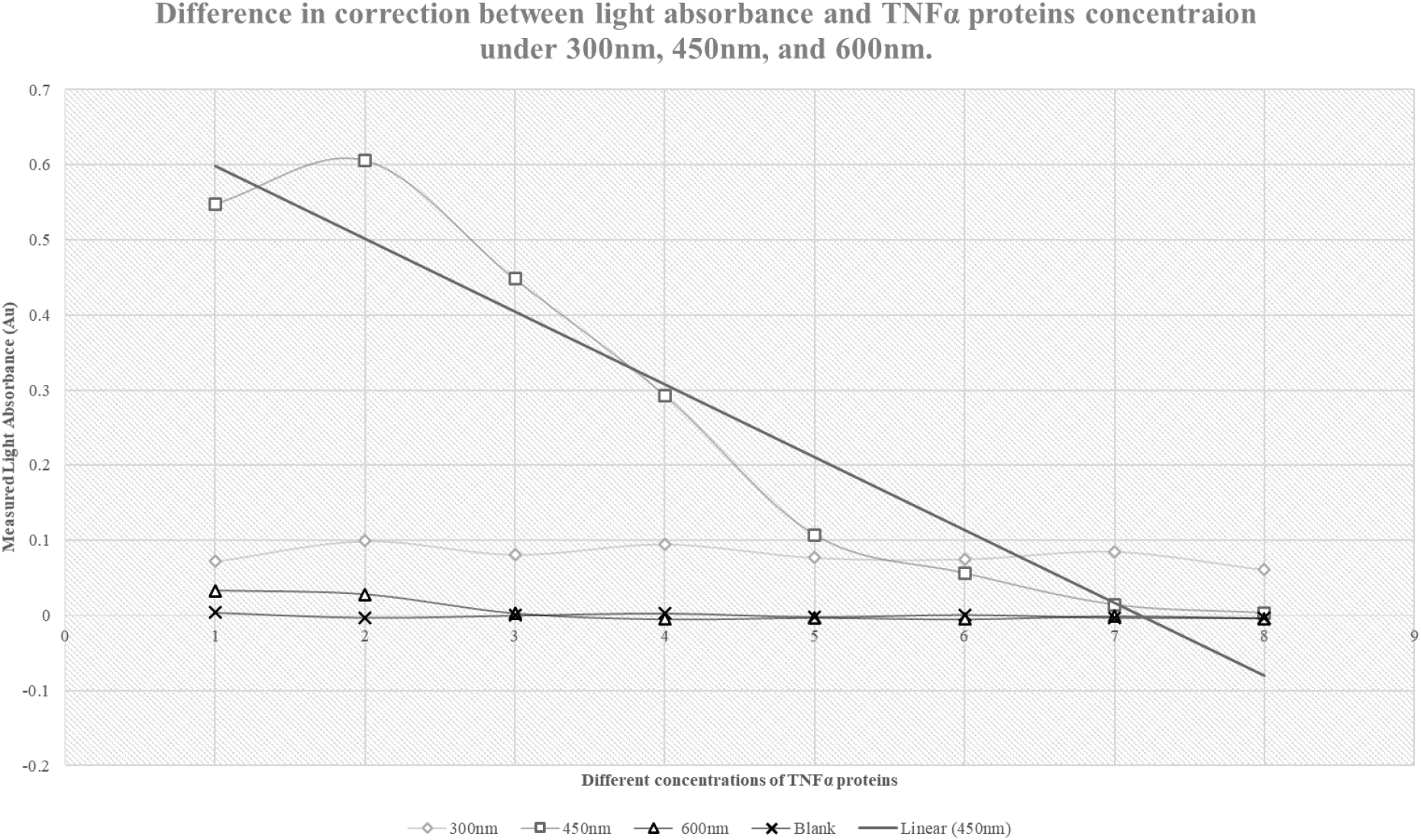
Difference in correlation between light absorbance and TNFα proteins concentration under 300 nm, 450 nm, and 600 nm: The data under 300 nm and 600 nm shows the slope near zero, which neither shows negative nor positive correlation between measured light absorbance and TNFα protein concentration. However, data under 450 nm shows strong negative correlation between measured light absorbance and TNFα proteins.

#### 3.2.3. Correlation between light Absorption and TNFα concentration

Under the same wavelength of 450nm, four columns including standard serum based ELISA result and salivary based ELISA in three different methods were analyzed using spectramax Pro with the Gen5 software. According to the data, the light absorbance level was greater in the wells with greater TNFα proteins: data Table 1 and 2. A well with 1000pg/mL TNFα protein showed maximum absorbance level of 0.601Au. in the standard ELISA, and 0.605Au. in the saliva based ELISA- Method 3. A well with 14.6pg/mL TNFα proteins showed maximum absorbance of 0.067Au in blood based ELISA, and 0.094Au in the saliva based ELISA- Method 3. Meanwhile, the data from chromogen blank and assay controls showed an absorbance level around 0.05 ± 0.01Au. Correlation between light absorption and TNFα concentration is drawn in Graph 2.

**Table. 1-.** Light absorbance for 8 different dilution samples in three different methods- Raw absorbance data.

| Column # | Column 1<br>(Standard) | Column 2 (Method 1) | Column 3 (Method 2) | Column 4 (Method 3) | Blank | Analysis<br>Condition |
| --- | --- | --- | --- | --- | --- | --- |
| A (1000pg/mL) | 0.601 | 0.166 | 0.055 | 0.605 | 0.057 | 450nm |
| B (500pg/mL) | 0.659 | 0.24 | 0.666 | 0.195 | 0.05 | 450nm |
| C (250pg/mL) | 0.501 | 0.178 | 0.413 | 0.135 | 0.053 | 450nm |
| D (125pg/mL) | 0.346 | 0.198 | 0.103 | 0.083 | 0.056 | 450nm |
| E (62.5pg/mL) | 0.16 | 0.055 | 0.117 | 0.061 | 0.051 | 450nm |
| F (31.2 pg/mL) | 0.109 | 0.057 | 0.089 | 0.066 | 0.054 | 450nm |
| G (15.6 pg/mL) | 0.067 | 0.061 | 0.065 | 0.094 | 0.05 | 450nm |
| H (0 pg/mL) | 0.056 | 0.079 | 0.068 | 0.105 | 0.05 | 450nm |

**Table. 2-.**
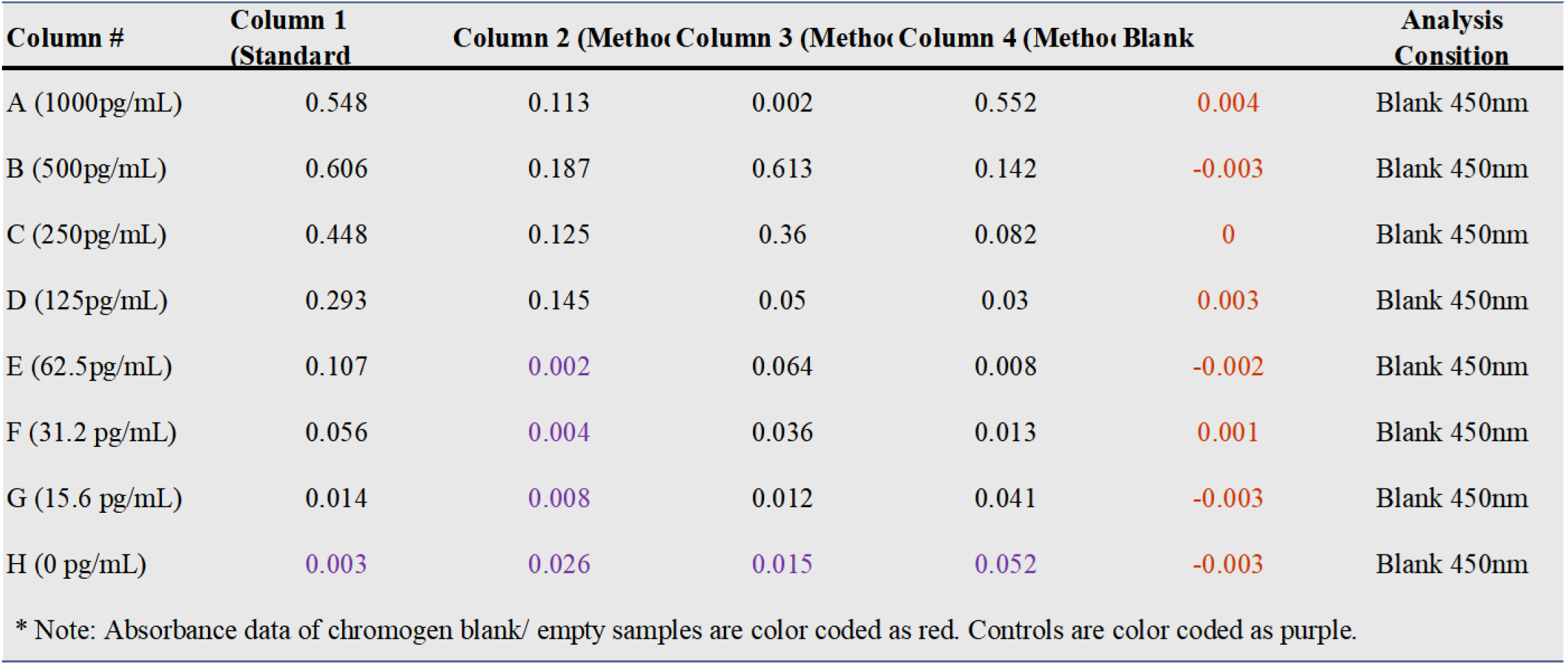
Light absorbance for 8 different dilution samples in three different methods- data with blank standardziation.

**Table. 3-.**
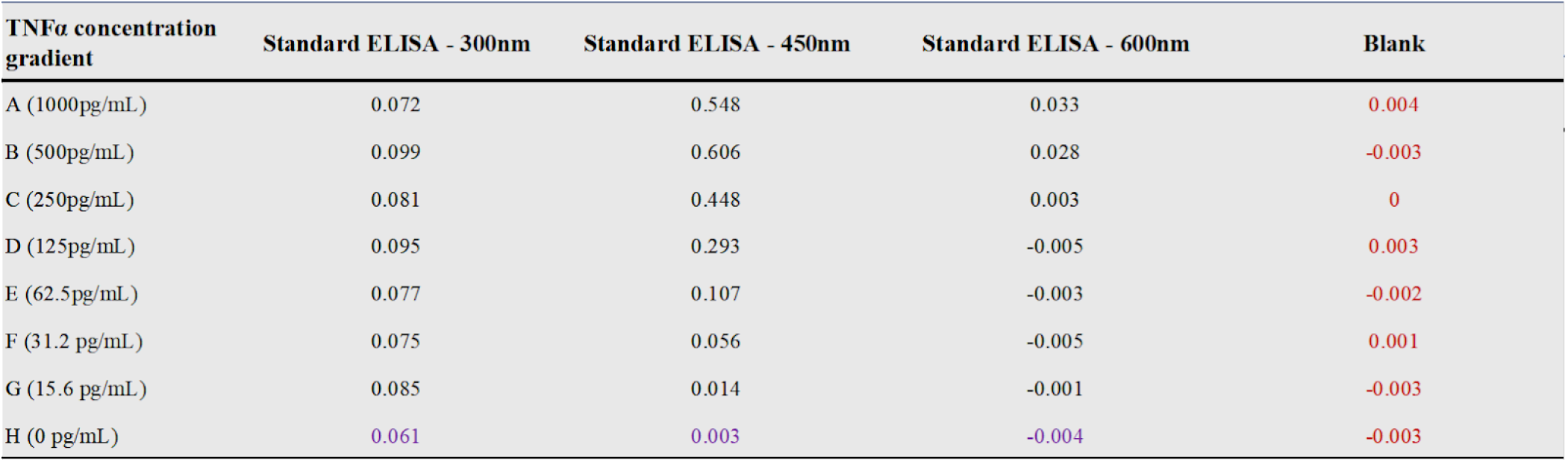
Different in correction between light absorbance and TNFa proteins concentraion under 300nm, 450nm, and 600nm. Measure of light absorbance (Au) of different TNFα diluted solutions analyzed under three different wavelengths: 300 nm, 450 nm, 600 nm: the graph of this data is shown below. *This data is color coded corresponding to Image 3.

**Graph. 2-.**
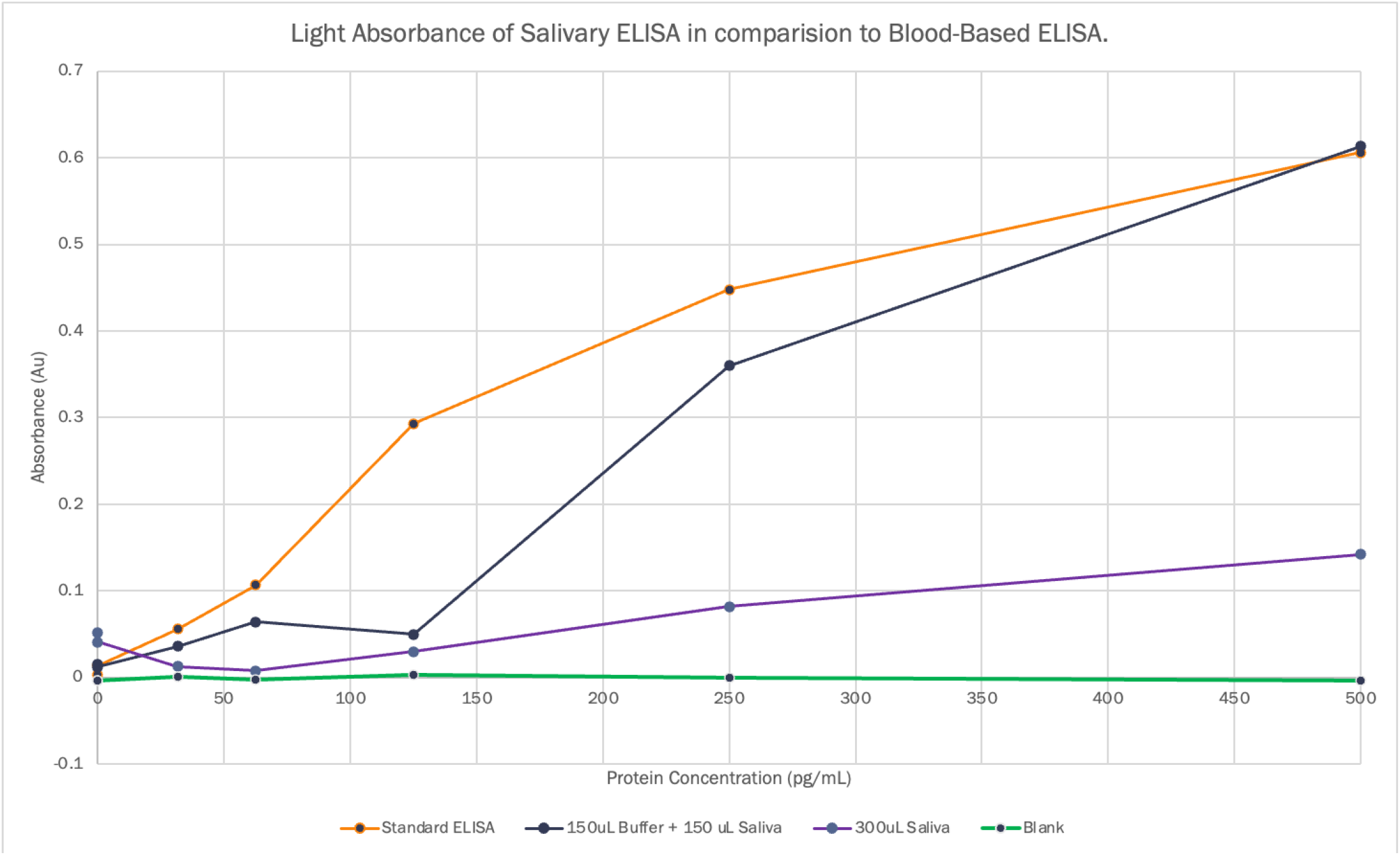
Measured light absorbance (Au) vs. different concentrations of diluted TNFα samples in three different groups: This graph shows the negative correlation between TNFα concentration and level of light absorbance. In this graph, the data is edited to mark the absorbance level of sample controls and chromogen blanks into 0.00 ± 0.01Au.

#### 3.2.4. Pearson Correlation between groups

In order to further analyze the efficiency of salivary based ELISA protocol, researchers analyzed the correlation using pearson correlation formula, which gives the correlation, association, and direction of relationship between two sets of datas:

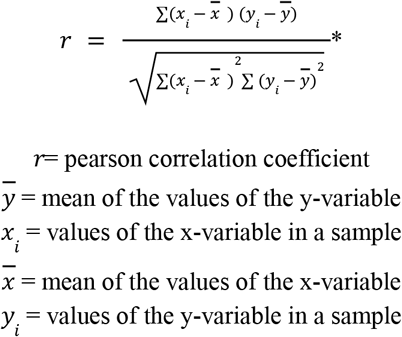

**\*The relations between pearson coefficient and relationship between sample groups are the following: 1**≥ *r***>0.7: a strong positive relationship, 0.7**≥*r***>0.5: a moderate positive relationship, 0.5r>-0.5: no relationship, -0.5r-0.7: a moderate positive relationship, and -0.7**≥*r*≥**1: a strong negative relationship**.

According to data from Table 1, the pearson coefficient between standard ELISA and saliva based ELISA -Method 2- showed 0.6686887643, and between standard ELISA and Method 3 showed 0.626571372. Therefore, researchers concluded that salivary based ELISA has moderate to high positive relationship with standard ELISA.

#### 3.2.5. Optical Density Analysis

Optical density of each sample wells were then analyzed in order to determine each radiant power that is passed through the material:

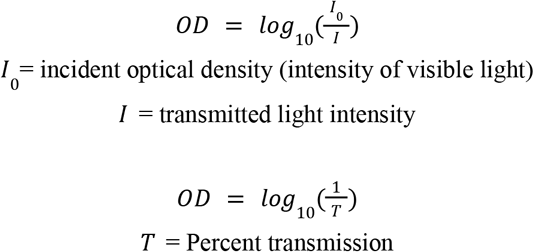

The calculated optical density is shown in Table 4 and drawn in Graph 3.

**Table. 4-.** Optical Density and transmission for standard blood serum based ELISA.

| Methods | absorbance | Transmission | Standard Optical Density | log (100/T) for Standard | log(100/T) |
| --- | --- | --- | --- | --- | --- |
| A (1000pg/mL) | 0.548 | 28.31391996 | 3.38 | 0.5289167 | 0.548 |
| B (500pg/mL) | 0.606 | 24.77422058 | 1.97 | 0.294466226 | 0.606 |
| C (250pg/mL) | 0.448 | 35.64511334 | 1.18 | 0.071882007 | 0.448 |
| D (125pg/mL) | 0.293 | 50.93308711 | 0.75 | -0.124938737 | 0.293 |
| E (62.5pg/mL) | 0.107 | 78.16278046 | 0.38 | -0.420216403 | 0.107 |
| F (31.2 pg/mL) | 0.056 | 87.90225168 | 0.22 | -0.657577319 | 0.056 |
| G (15.6 pg/mL) | 0.014 | 96.82778563 | 0.15 | -0.823908741 | 0.014 |
| H (0 pg/mL) | 0.003 | 99.31160484 | 0.08 | -1.096910013 | 0.003 |

**Graph. 3-.**
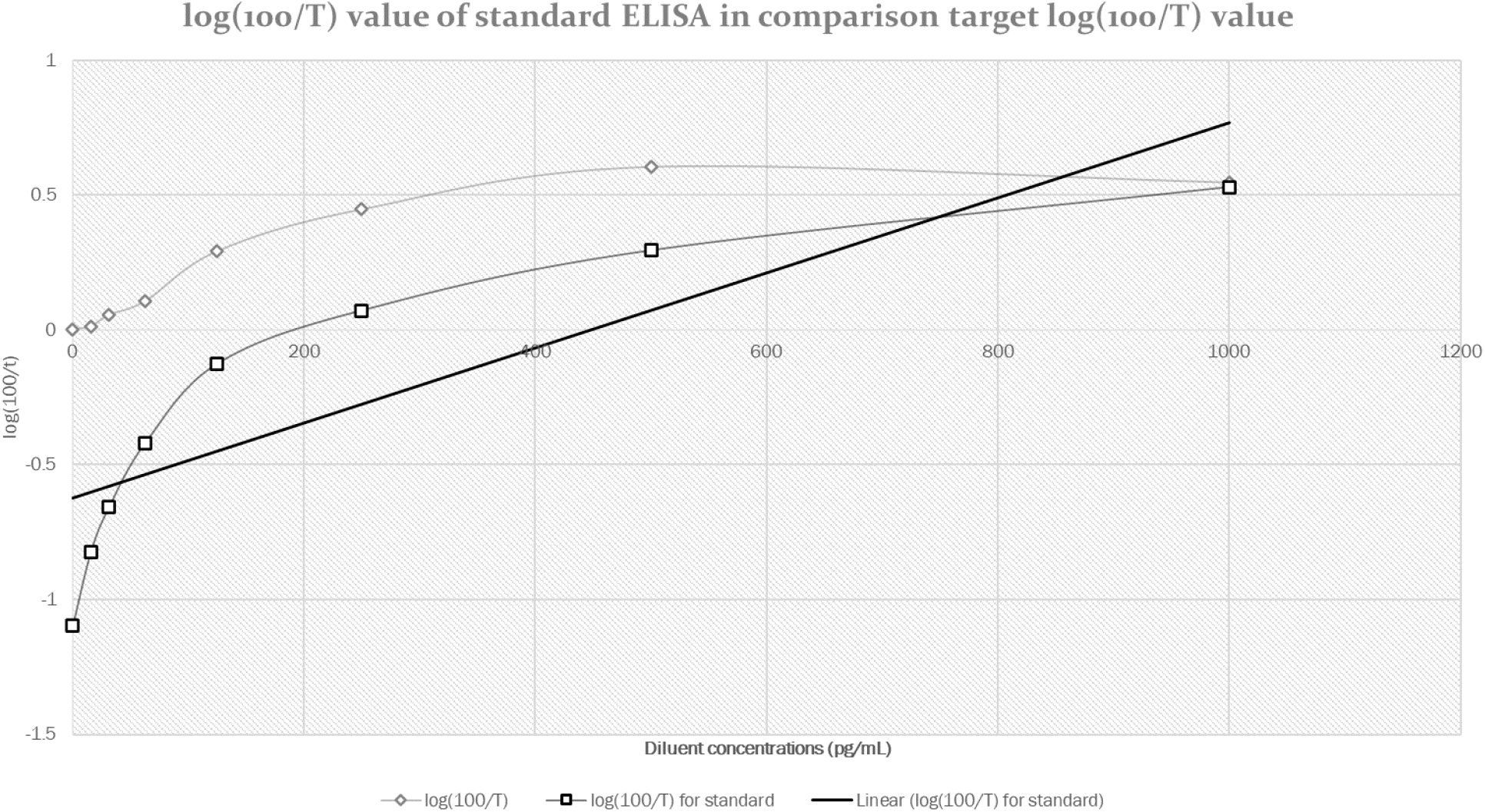
log(100/T) value of standard ELISA in comparison to target log(100/T) value.

## 4. Discussion

According to the data provided by Thermofisher, the range of the optical density calculated ranged from 0.08 to 3.38; therefore, the log(100/T) value ranged from -1.097 to 0.529. Meanwhile, the calculated log(100/T) value from the new salivary protocol ranged from 0.003 to 0.548. While the maximum value of log(100/T) in both protocols is 0.53 ± 0.01, the range of log(100/T) decreased in the new protocol from 1.626 to 0.545- data shown in data Table 5 and Graph 4.

**Table. 5-.** log(100/T) value from other research vs. calculated log(100/T) value from new salivary based ELISA protocol: Range of log(100/T) values of other research is 1.626, while range of log(100/T) value of new protocol is 0.545. The data is provided by thermofisher (*Thermofisher*).

| Table 5- log(100/T) datas from other research vs. calculated log(100/T) c |  |  |  |
| --- | --- | --- | --- |
| TNFα concentration | Standard Optical Density | log (100/T) for Standard | Calculated log(100/T) |
| 1000 | 3.38 | 0.5289167 | 0.548 |
| 500 | 1.97 | 0.294466226 | 0.606 |
| 250 | 1.18 | 0.071882007 | 0.448 |
| 125 | 0.75 | -0.124938737 | 0.293 |
| 62.5 | 0.38 | -0.420216403 | 0.107 |
| 31.2 | 0.22 | -0.657577319 | 0.056 |
| 15.6 | 0.15 | -0.823908741 | 0.014 |
| 0 | 0.08 | -1.096910013 | 0.003 |

Based on the data comparison, researchers concluded that results were different from datas from other research due to two big reasons. First of all, actual concentrations of TNFα in each well could have been a smaller concentration than expected due to the centrifuging and filtering process. Extravagant centrifuging and filtering processes could have filtered too much debris, even including TNFα proteins. Therefore, this could have caused the capturing antibodies harder to find the proteins to bind. Another hypothesis is due to the fact that the percentage of immunoglobulin for saliva is different from that of blood serum based. Percentage of IgG- immunoglobulin that is found in blood and other body fluids- antibody in sample is usually around 80%, while percentage of IgA- immunoglobulin that is found in respiratory tract, saliva, and tears- is around 13% ([Lakna](https://pediaa.com/what-is-the-difference-between-igg-igm-iga-ige-and-igd/)). Therefore, decrease in sensitivity could be due to difference in level of antibodies existing in samples. Nevertheless, researchers were able to conclude that the new salivary based ELISA protocol is capable of diagnosing inflammatory bowel disease as it can detect moderate amounts of TNFα proteins: the maximum value of log(100/T) was similar with data from other research and the range of values are wide enough- .003 to .548- to differentiate the samples into different diluent concentrations.

Even though researchers obtained the desired data, researchers concluded results still lacked validity due to lack of replicates that prevented obtaining more reliable statistical datas. Throughout the research, a total of 3 ELISA exams were performed in 5 different methods. In order to validify the data, more trials need to be performed in the future. Another shortcoming is the fact that each ELISA plate was analyzed at least a week after each experiment. The plates were stored in -80°C freezer after each experiment, and went through the thawing process before spectramax analysis. However, an analysis is supposed to be done within 2 hours of experiment completion in order to prevent the colors changing back to clear and proteins from degenerating. Hence, the process of freezing and thawing could have prevented researchers from getting more accurate results; for the first standard ELISA plate, especially, it went through multiple frequent freezing-thawing procedures. In fact, most of the proteins were degenerated and therefore were unable to proceed the analysis procedure.

Regardless of the shortcomings, a strength of this experiment is the fact that researchers were able to detect the color difference between the diluted TNFα saliva samples instead of blood serum samples. The range of the absorbance level from the salivary sample is wide enough to detect the different concentrations. Future work should include more trials of ELISA using salivary samples and analysis of the plate within 2 hours of experiment.

**Graph. 5-.**
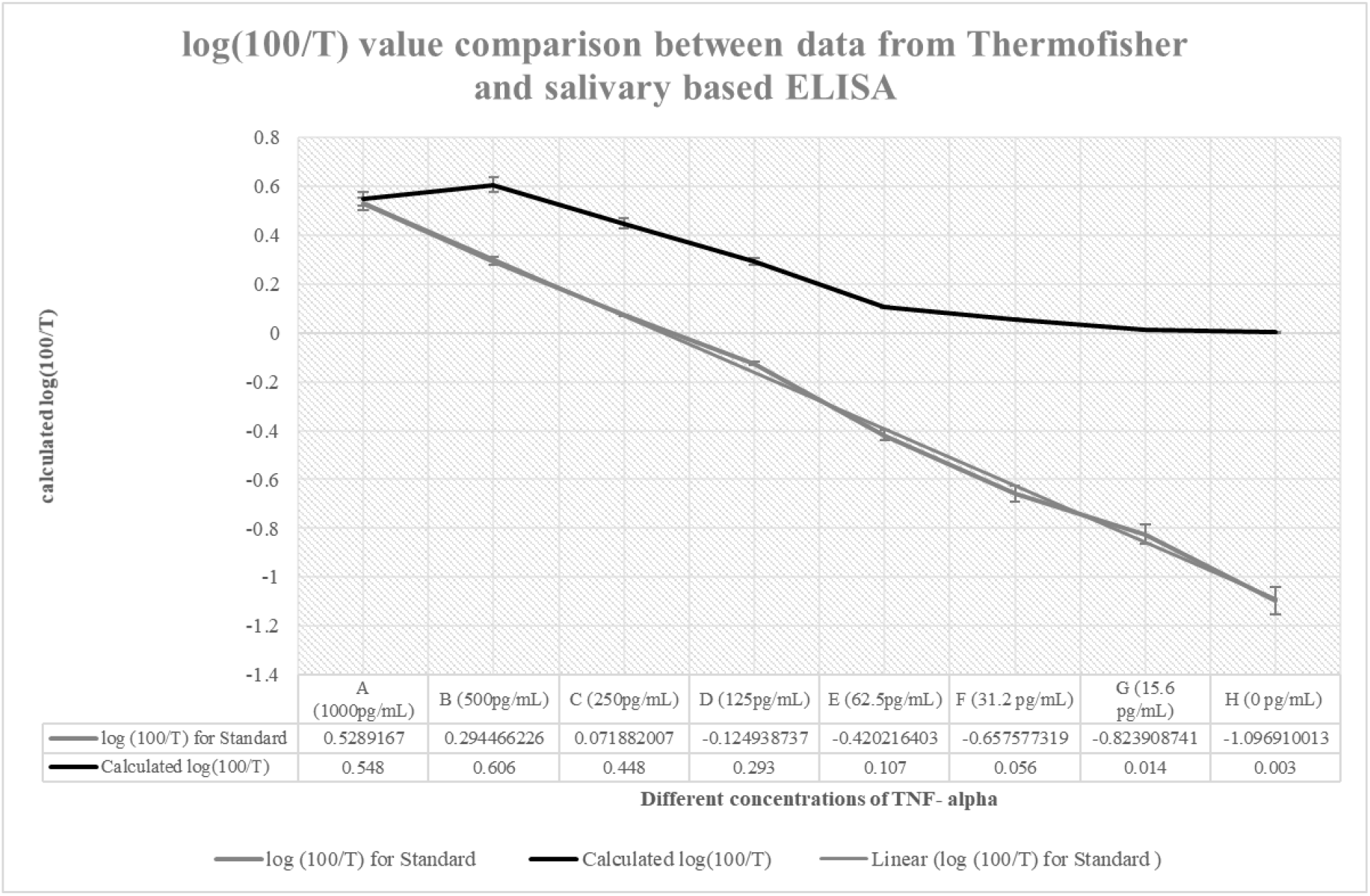
log(100/T) data comparison data given by *Thermofisher* to that of salivary based ELISA protocol: as shown on the graph above, the data range for measured log(100/T) is 1.081 less than data provided by *Thermofisher*. This means the sensitivity of the new protocol decreased, and it was less efficient to measure different concentrations of diluted TNFα.

## 5. Conclusion

- Data comparison between blood serum based ELISA and salivary based ELISA shows how detection of TNFα proteins is possible using salivary samples. The data correlation showed a moderate positive relation between blood and salivary ELISA, showing how each can be used interchangeably.
- Salivary ELISA lost 50% sensitivity from 31.2pg/mL to 15.6pg/mL TNFα concentration, but was able to differentiate diluted samples from 1000pg/mL to 31.2pg/mL TNFα concentration.
- More research needs to be done in order to validify the data and for more accurate results.
- Future directions for this project include modifying the saliva to buffer ratio in future trials to gain better recovery of protein with salivary input, as well as modifying the color analysis app to read multiple plates. Additionally, we would hopefully be able to modify the ELISA procedure to be an at-home diagnostic by reducing its run time and simplifying the sample collection procedure.

## Acknowledgements

The research was conducted with co-researcher Anjana Ganesh. It was also funded and supported by Timothy E. Riedel- research educator of DIY diagnostic stream under freshman research initiative at University of Texas at Austin- as well as funded by Bob and Cathy O’Rear. The reference data table was created by thermofisher (*[Thermofisher](https://assets.thermofisher.com/TFS-Assets/LSG/manuals/MAN0003931_KHC3011_HumanTNFalpha_ELISA_PI.pdf)*).

## References

Abigail L. Mandel, Hakan Ozdener & Virginia Utermohlen (2011) BRAIN-DERIVED NEUROTROPHIC FACTOR IN HUMAN SALIVA: ELISA OPTIMIZATION AND BIOLOGICAL CORRELATES, Journal of Immunoassay and Immunochemistry, 32:1, 18-30, DOI:

Amber J Tresca https://www.verywellhealth.com/is-there-a-cure-for-inflammatory-bowel-disease-1942489

Amber J. Tresca https://www.verywellhealth.com/is-crohns-autoimmune-5188106

Art Kushner, “Fake Saliva,” PF Online, Gardner Publications, Cincinnati, OH; http://www.pfonline.com/articles/clinics/1206cl_plate5.html (Last accessed 3/31/10).

Baumgart DC, Sandborn WJ (August 2012). “Crohn’s disease”. Lancet. 380 (9853): 1590–605. doi:10.1016/S0140-6736(12)60026-9. PMID 22914295

Brinkman, B. M., Zuijdeest, D., Kaijzel, E. L., Breedveld, F. C., & Verweij, C. L. (1995). Relevance of the tumor necrosis factor alpha (TNF alpha)-308 promoter polymorphism in TNF alpha gene regulation. Journal of inflammation, 46(1), 32–41.

Crohn’s Colitis Foundation, The Facts About Inflammatory Bowel Diseases. New York, NY: Crohn’s and Colitis Foundation of America; 2014. http://www.crohnscolitisfoundation.org/assets/pdfs/updatedibdfactbook.pdf [PDF-2.32MB]

Crohn’s Disease”. National Institute of Diabetes and Digestive and Kidney Diseases (NIDDK). Archived from the original on December 8, 2019. Retrieved December 8, 2019.

Hettegger, Peter et al. “High similarity of IgG antibody profiles in blood and saliva opens opportunities for saliva based serology.” PloS one vol. 14,6 e0218456. 20 Jun. 2019, doi:10.1371/journal.pone.021845610.1080/15321819.2011.538625

Hornbuckle, W. E., & Tennant, B. C. (1997). Gastrointestinal function. In Clinical biochemistry of domestic animals (pp. 367–406). Academic Press.

Howmuchisit.org https://www.howmuchisit.org/how-much-does-elisa-test-cost/

Jeffery M. Perkel https://www.biocompare.com/Editorial-Articles/137081-Microplate-Readers-Many-Options-for-Multiple-Applications/

Lakna. “What Is the Difference between IGG IGM IGA IGE and IgD.” Pediaa.Com, Pediaa, 14 Oct. 2019, https://pediaa.com/what-is-the-difference-between-igg-igm-iga-ige-and-igd/.

Loftus EV, Jr., Shivashankar R, Tremaine WJ, Harmsen WS, Zinsmeiseter AR. Updated Incidence and Prevalence of Crohn’s Disease and Ulcerative Colitis in Olmsted County, Minnesota (1970-2011). ACG 2014 Annual Scientific Meeting. October 2014.

MacMullan, M.A., Ibrayeva, A., Trettner, K. et al. ELISA detection of SARS-CoV-2 antibodies in saliva. Sci Rep 10, 20818 (2020). https://doi.org/10.1038/s41598-020-77555-4

Markham, R., Young, L., & Fraser, I. S. (1995). An amplified ELISA for human tumour necrosis factor alpha. European cytokine network, 6(1), 49–54.

Marks DJ, Rahman FZ, Sewell GW, Segal AW (February 2010). “Crohn’s disease: an immune deficiency state”. Clinical Reviews in Allergy & Immunology. 38 (1): 20–31. doi:10.1007/s12016-009-8133-2. PMC 4568313. PMID 19437144

Mayo Clinic: https://www.mayoclinic.org/diseases-conditions/inflammatory-bowel-disease/diagnosis-treatment/drc-20353320

MedlinePlus https://medlineplus.gov/ency/article/003332.htm

Morris, M. W., Stewart, S. A., Heisler, C., Sandborn, W. J., Loftus, E. V., Zello, G. A., … & Jones, J. L. (2018). Biomarker-based models outperform patient-reported scores in predicting endoscopic inflammatory disease activity. Inflammatory bowel diseases, 24(2), 277–285.

Nakamura, R. M., Matsutani, M., & Barry, M. (2003). Advances in clinical laboratory tests for inflammatory bowel disease. Clinica chimica acta, 335(1-2), 9–20.

Thermofisher. “Human TNF-a ELISA Kit Product Information Sheet.” Thermofisher, 19 Nov. 2019.

